# QUANTA: A quantum annealing framework for species tree estimation using weighted quartet amalgamation

**DOI:** 10.64898/2026.11.05.764558

**Authors:** Niloy Kumar Mondal, Md. Shamsuzzoha Bayzid

## Abstract

Reconstructing species trees from genome-scale data is challenging due to pervasive gene tree discordance. Maximizing quartet consistency is a widely used approach for species tree inference from multi-locus data. Quartet amalgamation approaches, such as Quartet Fiduccia–Mattheyse (QFM) and Quartet MaxCut (QMC) families, have emerged as highly accurate quartet-based species tree inference techniques, but their reliance on computationally expensive classical bipartitioning heuristics, such as Fiduccia–Mattheyses (FM) and MaxCut, limits their scalability. Here, we introduce QUANTA, a quantum annealing-based quartet amalgamation method that, to our knowledge, represents the first quantum formulation of quartet-based approaches and the first species tree estimation method for multi-locus data that runs on real quantum hardware. At the core of QUANTA is a novel, general formulation of weighted quartet bipartitioning as a quadratic unconstrained binary optimization (QUBO) problem that enables quantum annealing to replace the classical bipartitioning strategies underlying the QFM and QMC families of methods. We prove that our formulation uses the minimum possible number of qubits to represent bipartitions and that its energy function correctly orders satisfied, deferred, and violated quartets for any positive penalty weights. We implemented QUANTA on real D-Wave quantum annealing hardware, and evaluated it on a wide range of simulated and biological datasets. QUANTA achieves accuracy comparable to the state-of-the-art methods, including ASTRAL and wQFM, under challenging evolutionary conditions. We also reanalyzed several biological datasets, including the avian phylogenomic dataset comprising 14,446 genes across 48 genomes, on which QUANTA reconstructed a species tree with a higher quartet score than wQFM. Remarkably, despite running on current noisy intermediate-scale quantum (NISQ) hardware, QUANTA’s bipartition technique achieved 4100× speedup over the FM bipartitioning technique in wQFM on bipartition containing 90–200 taxa, while maintaining comparable tree accuracy. By establishing a theoretically grounded, hardware-independent quantum optimization framework for weighted quartet amalgamation, QUANTA opens a new direction for quantum-assisted phylogenomics and provides a foundation for increasingly scalable and accurate species tree estimation as quantum computing hardware advances. QUANTA is publicly available at https://github.com/QuitttCat/QUANTA-Phylogenomics.

## 1 Introduction

Modern phylogenomic studies infer species trees from hundreds or thousands of genes, with molecular sequence data typically partitioned into recombination-free genomic regions, referred to as loci or genes. Different genes often have conflicting evolutionary histories due to biological processes such as incomplete lineage sorting (ILS) [1], which is modeled by the multispecies coalescent (MSC) [2, 3]. In the presence of gene tree discordance, traditional approaches such as concatenation, which combines gene alignments into a single supermatrix to infer a species tree, can be statistically inconsistent [4, 5] and may produce incorrect trees with high support [6]. Consequently, summary methods, which first estimate individual gene trees and then combine them to infer a species tree, have become increasingly popular [7]. Many of these methods are statistically consistent under the MSC model [8–20].

Over the past decade, quartet-based methods have emerged as some of the most widely used approaches to species tree estimation, owing to their theoretical guarantees and high accuracy. ASTRAL, the most popular summary method, seeks to maximize the total number of quartet trees shared between the species tree and the input gene trees. An alternative approach is quartet amalgamation, which first infers evolutionary relationships among subsets of four taxa and then combines these quartets into a single species tree [16, 21–27]. Quartet amalgamation methods, such as the QFM and QMC families, have attracted considerable interest because quartets can be inferred accurately from various sources of evolutionary information and subsequently combined to reconstruct a species tree. Weighted QFM (wQFM) was shown to be more accurate than ASTRAL on challenging model conditions characterized by high ILS and high GTEE [28, 29]. Unlike summary methods that rely on estimated gene trees and can be sensitive to gene tree estimation errors (GTEEs) [7, 30, 31], quartet amalgamation can also be applied to quartets inferred directly from sequence data. For example, SVDquartets [16] uses techniques from algebraic statistics to infer quartet relationships directly from sequence alignments, which can then be amalgamated into a species tree. This flexibility, together with the high accuracy of quartet amalgamation methods, makes them an attractive approach to species tree estimation. However, these amalgamation methods are not practical for large-scale phylogenomics as there are *θ*(*n*^4^) quartets and the number of candidate trees (2*n* − 5)!! grows super-exponentially with the number of taxa [32].

Here, we explore quantum computing as an alternative approach to improving the scalability and accuracy of species tree estimation. Quantum computing offers potential advantages over classical methods by considering many candidate solutions at once through superposition and entanglement to explore large solution spaces, and by escaping locally optimal solutions through quantum tunneling [33–35]. Moving beyond textbook quantum algorithms, quantum computing is now becoming practical: companies such as IBM [36], Google [37] and D-Wave [38] build quantum processors that can be accessed remotely through the cloud. These processors, however, belong to the noisy intermediate-scale quantum (NISQ) era [39]: their limited numbers of qubits and their susceptibility to noise restrict what they can currently achieve. The full potential of quantum computing for scalability and accuracy is expected to be realized as hardware advances toward fault-tolerant quantum computers. Computational biology has begun to explore this paradigm, with proof-of-concept quantum approaches to protein folding [40], genome assembly [41], RNA folding [42] and ligand pose generation [43]. Phylogenetics is particularly well suited to this exploration, since its central problems require searching a super-exponentially large space of trees, yet work in this direction has remained largely theoretical. Existing studies have explored graph-cut tree reconstruction [44–46], maximum parsimony [47] and tree containment [48], mostly with quantum-inspired, classical or simulated solvers, and only as proofs of concept on real hardware for trees of at most 29 taxa [46]. None of these studies, however, has addressed species tree estimation from multi-locus data, or run phylogenetic inference on real quantum hardware beyond a few dozen taxa.

In this study, we take a step toward filling this gap and present **QU**antum **AN**nealing for **T**ree **A**malgamation (QUANTA), which, to the best of our knowledge, is the first quantum formulation for species tree inference from multi-locus data. Unlike existing quartet amalgamation methods, which employ method-specific classical heuristics such as FM and MaxCut to identify bipartitions, QUANTA provides a novel, general-purpose quantum-compatible optimization formulation of weighted quartet bipartitioning. This formulation is independent of the particular amalgamation framework and quantum hardware and can therefore be integrated into different quartet-based reconstruction pipelines, including those of the QFM and QMC families. We formulate this step as a quadratic unconstrained binary optimization (QUBO) [49] problem in a resource efficient way. QUANTA uses the minimum possible number of binary variables to encode bipartitions. We solved the formulated QUBO problems on D-Wave quantum computers through D-Wave’s cloud service [38]. We instantiate QUANTA within the amalgamation framework of wQFM to demonstrate its practical advantages. Our experiments show that QUANTA achieves accuracy comparable to that of state-of-the-art methods, including wQFM and ASTRAL, while solving bipartitions of 90–200 taxa about 4100 times faster than the FM heuristic of wQFM. On the challenging avian dataset [50] with 14,446 genes sampled from 48 bird genomes, QUANTA reconstructs a species tree with a higher quartet score than wQFM. Moreover, our formulation offers a promising starting point for extending quantum optimization to the more general quatet-based optimization problem addressed by methods such as ASTRAL. Thus, QUANTA lays the foundation for a broader quantum framework for quartet-based species tree estimation, with the potential to greater scalability and improved phylogenetic accuracy as quantum computing hardware advances.

## 2 Materials and Methods

### 2.1 Problem Definition: Quartet Amalgmation

We address the *weighted Maximum Quartet Consistency* (wMQC) problem introduced by QMC and QFM families of quartet amalgamation methods [25, 26, 28].

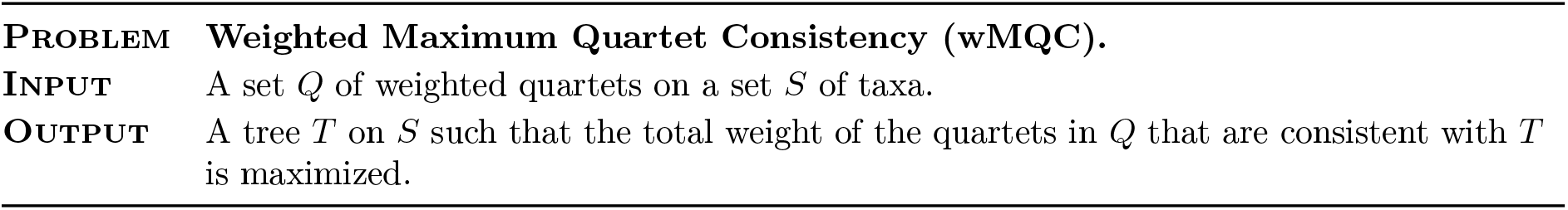

| **Problem** | **Weighted Maximum Quartet Consistency (wMQC).** |
|---|---|
| **Input** | A set *Q* of weighted quartets on a set *S* of taxa. |
| **Output** | A tree *T* on *S* such that the total weight of the quartets in *Q* that are consistent with *T* is maximized. |

Quartet amalgamation methods, such as the QFM and QMC families, follow a divide-and-conquer approach to reconstruct a species tree from a set of input quartets. They first partition the taxa into two disjoint subsets by seeking a bipartition that maximizes agreement with the input quartets. For a given bipartition, a quartet is considered *satisfied* if its two pairs of taxa lie on opposite sides, *violated* if the bipartition separates the taxa in a conflicting manner, and *deferred* if it remains unresolved by the bipartition. A visualization of the quartet satisfaction, violation, and deferral criteria is provided in Supplementary Section 1 (Fig. S1). Candidate bipartitions are evaluated by rewarding satisfied quartets and penalizing violated ones, while deferred quartets are retained for subsequent recursive steps. The partitioning process is then applied recursively to each subset, with artificial taxa introduced to preserve the relationships between the subsets. Once the recursive steps are completed, the resulting subtrees are amalgamated, using the dummy taxa, to obtain a species tree containing all input taxa. QFM and QMC share this framework but differ in their bipartitioning strategies: QFM uses the Fiduccia–Mattheyses heuristic, while QMC employs MaxCut. In this work, we replace these classical bipartitioning strategies with a quantum optimization framework based on a quadratic unconstrained binary optimization (QUBO) formulation.

### 2.2 QUBO Formulation of Weighted Quartet Bipartitioning Problem

Many combinatorial optimization problems involve binary decisions, such as assigning each taxon to one of two groups, with the goal of minimizing an overall cost. Quadratic Unconstrained Binary Optimization (QUBO) is a general way to express such problems [49]. Using QUBO therefore takes two steps: encode the problem as a matrix of these costs, and pass the matrix to a solver that returns low-cost assignments. The quantum computers directly takes this QUBO matrix as input and return yes/no assignment for binary decision variables.

Formally, let *Q* be an *n* × *n* upper-triangular matrix of real coefficients. A QUBO problem seeks a binary vector *x* = (*x*_1_, …, *x*_*n*_) ∈ {0, 1}^*n*^ that minimizes the objective, often called the energy,

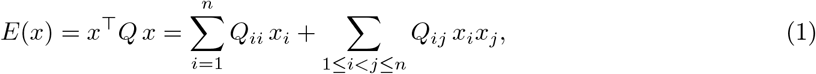

where the Q encodes of the cost function. So, to apply QUBO for our problem we need two design decisions, one for encoding binary decision and the other for constructing cost matrix. In the following, we discuss our design decision to convert Weighted quartet bipartitioning problem into QUBO problem in details with validation and proof.

#### Binary Decision Varible Encoding

In wQFM, each division step splits the current set of taxa *P* into two subsets *P*_*a*_ and *P*_*b*_, so the natural decision is, for each taxon, on which side of the split it goes.

We therefore associate each taxon *i* ∈ *P* with a binary decision variable *x*_*i*_, where *x*_*i*_ = 0 if *i* ∈ *P*_*a*_ and *x*_*i*_ = 1 if *i* ∈ *P*_*b*_, so that the vector *x* ∈ {0, 1}^|*P* |^ encodes a complete bipartition (*P*_*a*_ | *P*_*b*_).

On a quantum computer, each binary decision variable is represented by one logical qubit, the basic unit of quantum information. Unlike a classical bit, a qubit can exploit superposition, a phenomenon of quantum mechanics in which a system exists in a weighted combination of several states at once; when measured, the qubit collapses to 0 or 1 with probabilities given by these weights [34]. Current quantum computers, however, offer only a limited number of qubits [39], so the size of the problems they can solve depends directly on how many variables an encoding requires. Our encoding uses only one variable per taxon and, with a fixed reference taxon, attains the minimum possible (Theorem 2.1); it therefore makes the most of the available qubits and allows larger subproblems to be solved on today’s hardware.

##### Theorem 2.1

(Minimal Encoding Bound). *Let P be a set of n* ≥ 3 *taxa. Any encoding that represents every bipartition of P by a distinct assignment of m binary variables requires m* ≥ *n* − 1.

*Proof*. A bipartition (*P*_*a*_ | *P*_*b*_) of *P* is an unordered pair of nonempty disjoint subsets whose union is *P* . Every subset *S* ⊆ *P* other than ∅ and *P* defines one, with *P*_*a*_ = *S*, and *S* and *P* \ *S* define the same bipartition. Hence *P* has (2^*n*^ − 2)*/*2 = 2^*n*−1^ − 1 bipartitions. Since *m* binary variables take at most 2^*m*^ distinct values, representing all bipartitions requires 2^*m*^ ≥ 2^*n*−1^ − 1. For *n* ≥ 3 we have 2^*n*−1^ − 1 *>* 2^*n*−2^, and therefore *m* ≥ *n* − 1.

Once a reference taxon is fixed, our encoding meets this bound exactly. With one variable per taxon, an assignment *x* and its complement 1 − *x* describe the same bipartition with the two sides swapped, so the *n* variables carry one redundant degree of freedom. Fixing any one taxon to either side removes this redundancy. Indexing the taxa of *P* arbitrarily as 1, …, *n*, we set *x*_1_ = 0 without loss of generality; the index carries no meaning, and any taxon would serve equally well. The remaining *n* − 1 variables then determine *P*_*b*_ ⊆ *P* \ {1}, and each nonempty choice of *P*_*b*_ yields a different bipartition. Every bipartition of *P* is therefore represented exactly once using *n* − 1 variables, which matches the lower bound of Theorem 2.1, so no binary encoding can use fewer.

#### Cost Matrix Construction

We have designed the encoding part, we now explain our second design decision following the encoding method: the construction of the cost matrix *Q*. Our goal is to reconstruct a phylogenetic tree that maximizes the total weight of satisfied quartets. At each division step, a quartet whose taxa fall on both sides of the bipartition is in one of three states [26]. For a quartet *q* = ((*a, b*), (*c, d*)), the state of *q* is

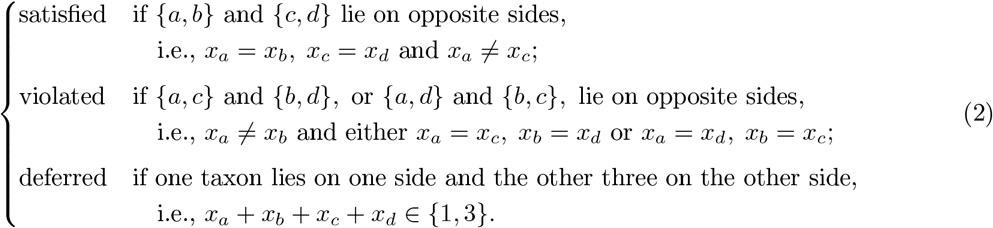

A deferred quartet remains unresolved at this step and is left for later steps of the recursion. Figure 1 in the supplementary material illustrates the three states.

**Figure 1:**
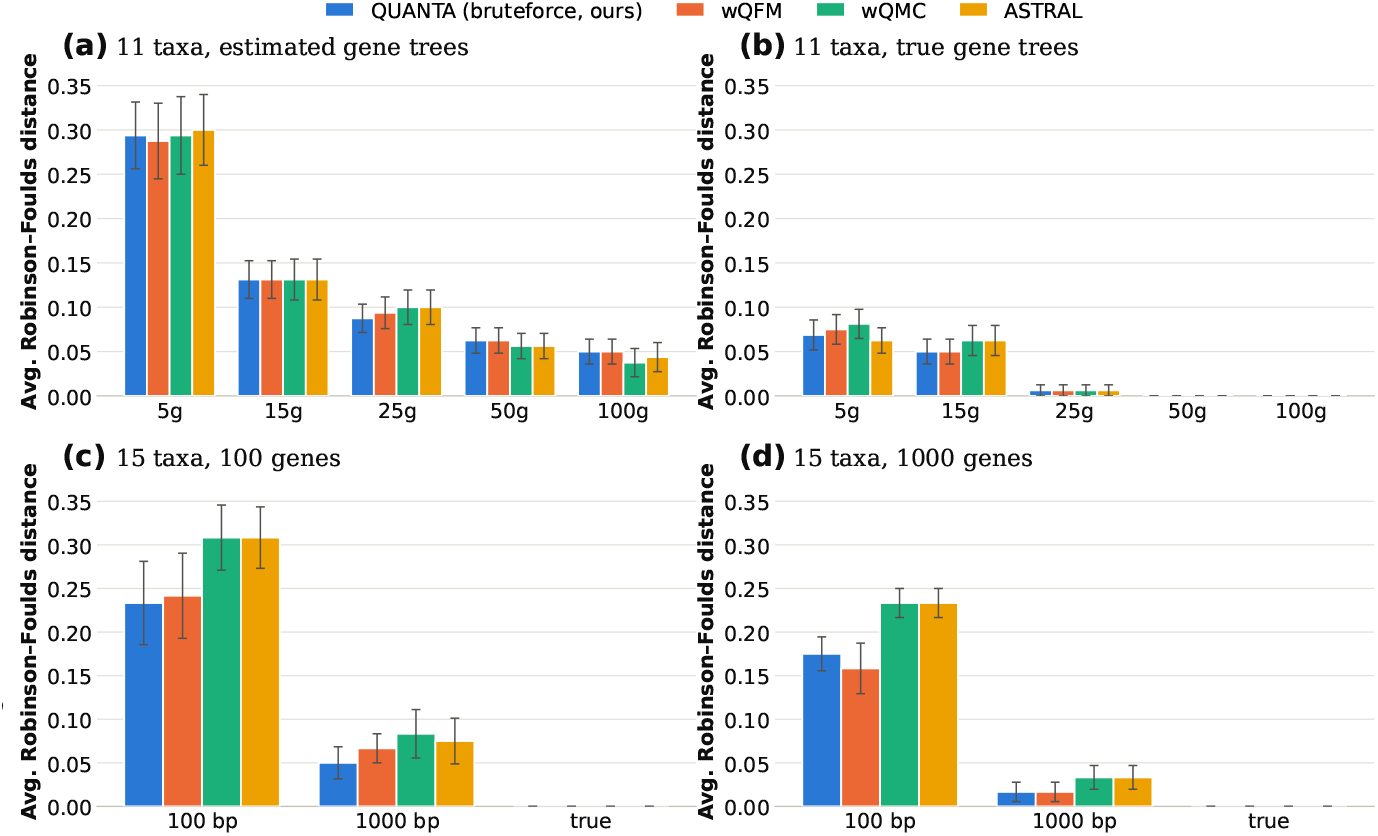
Average RF rates on the 11-taxon dataset with (a) estimated and (b) true gene trees, averaged over 20 replicates, and on the 15-taxon dataset with (c) 100 and (d) 1000 genes, averaged over 10 replicates. Error bars show standard errors.

We now construct the cost matrix, denoted by *Q*. Our goal is that satisfied quartets receive the lowest cost, violated quartets the highest, and deferred quartets a cost in between. Consider a quartet *q* = ((*a, b*), (*c, d*)) with weight *w*_*q*_ *>* 0. To track whether a bipartition satisfies *q*, we define the *satisfaction indicator χ*_*q*_ : {0, 1}^*n*^ → {0, 1}, which equals 1 if the assignment *x* satisfies *q* and 0 otherwise. Using the Iverson bracket [*P* ], which equals 1 if the statement *P* holds and 0 otherwise, it can be written as

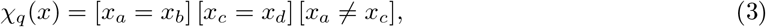

since *q* is satisfied exactly when each pair stays together and the two pairs lie apart. Ideally, the cost of a bipartition would then simply be − Σ_*q*∈Q_ *w*_*q*_ *χ*_*q*_(*x*), the negated total weight of the quartets it satisfies, where Q is the set of weighted quartets. However, expressing *χ*_*q*_ as a polynomial in *x* yields a fourth-degree expression

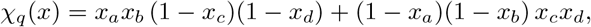

which contains the quartic term 2*x*_*a*_*x*_*b*_*x*_*c*_*x*_*d*_. A QUBO admits only linear and pairwise (quadratic) terms, so it cannot express this cost directly. Reducing the quartic term to quadratic form would require auxiliary variables [51], which would cost extra qubits and break the minimal encoding of Theorem 2.1.

We therefore follow the design of the cost function of Reaz et al. [26], also adopted by wQFM [28], which scores a bipartition by rewarding satisfied quartets and penalizing violated ones at the same time. Instead of rewarding a satisfied quartet as a whole, we penalize the mistakes that keep a quartet from being satisfied. There are two kinds of mistakes: splitting a pair that should stay together (*a* from *b*, or *c* from *d*), and placing two taxa from different pairs on the same side (one of *a, b* with one of *c, d*). We count the two kinds of mistakes with Iverson brackets:

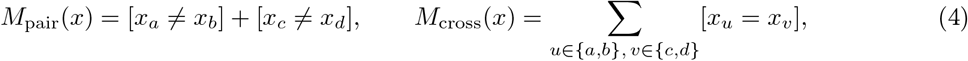

where *M*_pair_ is the number of split pairs and *M*_cross_ the number of cross pairs that share a side. A quartet is satisfied exactly when it makes no mistake: *M*_pair_(*x*) = 0 forces *x*_*a*_ = *x*_*b*_ and *x*_*c*_ = *x*_*d*_, and given this, *M*_cross_(*x*) = 0 holds exactly when *x*_*a*_ ≠ *x*_*c*_. Each bracket compares only two taxa and is therefore quadratic. For *x*_*i*_, *x*_*j*_ ∈ {0, 1}, the expression *x*_*i*_ − *x*_*j*_ ∈ {−1, 0, 1} vanishes exactly when *x*_*i*_ = *x*_*j*_, and *x*_*i*_ + *x*_*j*_ − 1 ∈ {−1, 0, 1} vanishes exactly when *x*_*i*_ ≠ *x*_*j*_; squaring them and using *x*^2^ = *x*_*i*_ gives

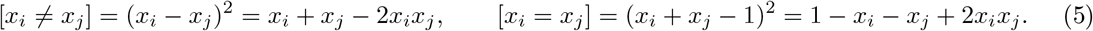

We penalize each mistake and scale the penalty by the weight of the quartet, which gives the cost

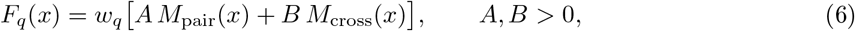

where *A* is the penalty for a split pair and *B* the penalty for a cross pair sharing a side. Substituting Equation (5) into Equation (4), and expanding Equation (6) we get,

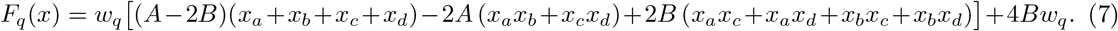

The constant 4*Bw*_*q*_ does not depend on *x*, so we drop it and obtain the energy of *q, E*_*q*_(*x*) = *F*_*q*_(*x*) − 4*Bw*_*q*_. The energy of a bipartition is the sum over all quartets, *E*(*x*) = Σ_*q*∈Q_ *E*_*q*_(*x*), which contains only linear and quadratic terms and therefore has exactly the form of a QUBO.

A QUBO is fully determined by its matrix, since its energy is *E*(*x*) = *x*^⊤^*Q x* (Equation (1)). To build *Q*, we therefore need only two kinds of numbers from *E*(*x*): the coefficient of each *x*_*i*_, which goes on the diagonal as *Q*_*ii*_, and the coefficient of each product *x*_*i*_*x*_*j*_, which goes above the diagonal as *Q*_*ij*_ with *i < j*. Both come directly from the expansion of *E*_*q*_. Every quartet adds (*A* − 2*B*) *w*_*q*_ for each of its four taxa, −2*A w*_*q*_ for each of its two pairs, and 2*B w*_*q*_ for each of its four cross pairs. Each entry of *Q* is therefore just a sum of quartet weights, grouped by the role the taxa play: *ω*_*i*_ = Σ_*q*∈Q_ *w*_*q*_ [*i* ∈ *q*] is the weight of the quartets containing 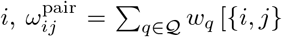 is a pair of *q*] the weight of those that pair *i* with *j*, and 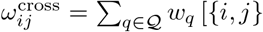 is a cross pair of *q*] the weight of those that place *i* and *j* in different pairs. This gives

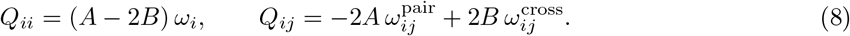

The result is easy to interpret. Quartets that pair *i* with *j* make *Q*_*ij*_ more negative and, in effect, pull the two taxa to the same side, while quartets that place them in different pairs make *Q*_*ij*_ more positive and push them apart. By minimizing *x*^⊤^*Q x*, the solver weighs all these pulls and pushes at once and seeks the bipartition that agrees best with the quartets. We have thus obtained the cost matrix *Q* that we set out to design, completing the QUBO construction. *Q* ranks the quartet states as intended (Proposition 2.2).

##### Proposition 2.2

(Quartet ordering). *Let q* = ((*a, b*), (*c, d*)) *be a quartet with weight w*_*q*_ *>* 0. *For any penalties A, B >* 0, *the QUBO matrix Q assigns q the energy*

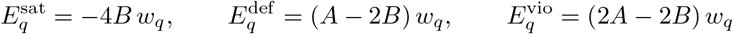

*when q is satisfied, deferred and violated, respectively; hence* 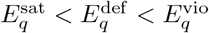.

*Proof*. By Equations (6) and (7), for any *x* ∈ {0, 1}^*n*^,

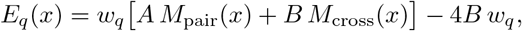

so it suffices to evaluate *M*_pair_ and *M*_cross_ in each state.

<u>Satisfied</u> (*ab* | *cd*): *x*_*a*_ = *x*_*b*_, *x*_*c*_ = *x*_*d*_ and *x*_*a*_ ≠ *x*_*c*_. Both pair indicators and all four cross indicators vanish, so (*M*_pair_, *M*_cross_) = (0, 0) and 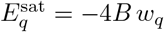.

<u>Deferred</u> (three taxa on one side): let *ℓ* be the lone taxon, *p* its partner in *q*, and {*u, v*} the other pair, so that *x*_*ℓ*_ ≠ *x*_*p*_ and *x*_*u*_ = *x*_*v*_ = *x*_*p*_. Then *M*_pair_ = [*x*_*ℓ*_ ≠ *x*_*p*_] + [*x*_*u*_ ≠ *x*_*v*_] = 1 and *M*_cross_ = [*x*_*ℓ*_ = *x*_*u*_] + [*x*_*ℓ*_ = *x*_*v*_] + [*x*_*p*_ = *x*_*u*_] + [*x*_*p*_ = *x*_*v*_] = 2, so 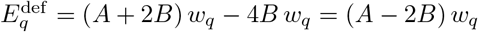.

<u>Violated:</u> the swap *c* ↔ *d* leaves both counts unchanged, so we may assume the split *ac* | *bd*, i.e., *x*_*a*_ = *x*_*c*_ ≠ *x*_*b*_ = *x*_*d*_. Then *M*_pair_ = [*x*_*a*_ ≠ *x*_*b*_] + [*x*_*c*_ ≠ *x*_*d*_] = 2 and *M*_cross_ = [*x*_*a*_ = *x*_*c*_] + [*x*_*a*_ = *x*_*d*_] + [*x*_*b*_ = *x*_*c*_] + [*x*_*b*_ = *x*_*d*_] = 2, so 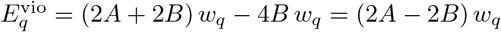.

Finally, 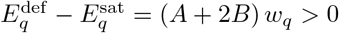 and 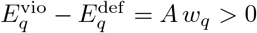 since *A, B, w*_*q*_ *>* 0.

If all four taxa of *q* fall on the same side, then *E*_*q*_(*x*) = 0, so the trivial bipartition with all taxa on one side also has energy 0; such degenerate solutions can be excluded by an optional balance penalty (Supplementary Section 1), and in practice we discard any sample with fewer than two taxa on either side and take the next lowest-energy sample.

This completes the construction of *Q*. Because *Q* depends only on the weighted quartets, it is solver-agnostic: the same matrix can be passed unchanged to a quantum computer, a classical heuristic, or an exact search. With the goal of finding most accurate bipartitions as fast as possible, we solve it by quantum annealing, which builds on the principles of adiabatic quantum computing [52, 53]. Unlike the Fiduccia–Mattheyses (FM) heuristic of QFM [26] and wQFM [28], which refines a bipartition by moving one taxon at a time, quantum annealing starts from a superposition of all 2^*n*^ bipartitions and gradually concentrates probability on low-cost ones [52, 54], while quantum tunnelling lets it pass through energy barriers that trap local search [53] (more details in Supplementary Section 3).

## 3 Experimental Studies

We evaluated QUANTA on a wide range of previously used simulated and biological datasets.

### Simulated Datasets

We used two biologically-based simulated datasets (the 37-taxon mammalian and 48-taxon avian simulated datasets) studied in [55]. We also used 11-taxon, 15-taxon, 101-taxon, and 200-taxon datasets from prior studies [55–57].

### Biological Datasets

We analyzed three biological datasets: the avian dataset comprising 48 taxa and 14,446 genes [50], the 37-taxon mammalian dataset from Song *et al*. [58], and the amniota dataset from Chiari *et al*. [59].

### Methods compared

We compared QUANTA with two leading weighted quartet amalgamation methods, wQFM, wQMC, and the most popular quartet-based summary method ASTRAL (ASTER v1.25.4.8 [60]). Moreover, to separate the contribution of the formulation from that of the quantum hardware, we solved the same QUBOs with two classical solvers, simulated annealing [61] and tabu search [62], instead of the quantum annealer. We compared the estimated trees (on simulated datasets) with the model species tree using normalized Robinson-Foulds (RF) distance [63] to measure the tree error.

### Bipartition Solver Details

We solved the QUBOs on D-Wave quantum annealers through D-Wave’s cloud service [38]. In our formulation, each taxon is one QUBO variable, that is, one logical qubit on the quantum processor (QPU). Current D-Wave systems can solve such QUBOs either directly on the QPU or with a hybrid quantum–classical solver, which divides the problem between classical processors and the QPU. Previous experiments on similar optimization problems [64, 65] and our estimates based on D-Wave’s qubit layouts and documented solver limits indicate that the QPU can solve dense QUBOs of up to about 175–184 variables, and that the hybrid solver can accept dense QUBOs of up to about 20,000 variables. Since quantum computing access is costly and limited, within our QPU cost and time budget, we experiment upto 200 taxa problems. These sizes reflect our QPU budget, not a hardware limit: because each taxon is one variable, the same QUBO with the hybrid solver could handle partitions of up to about 20,000 taxa (details in Supplementary Section 5). We chose a safe threshold: subproblems with up to 150 taxa are solved directly on the QPU, and larger ones are sent to the Leap hybrid solver. We also reduced QPU usage by solving partitions of at most 20 taxa exactly on a CPU by exhaustive (brute-force) search, since checking all 2^20^ ≈ 10^6^ assignments takes only seconds and guarantees the optimal solution. To assess the contribution of the quantum hardware, we also solved the same QUBOs with two classical heuristics, simulated annealing [61] and tabu search [62], using the open-source implementations in D-Wave’s `dwave-samplers` package (v1.8.0) on a standard CPU.

### 3.1 Results on simulated datasets

#### 11- and 15-taxon datasets

On these small simulated datasets, every partition has at most 20 taxa, so QUANTA solves each division step exactly by exhaustive search (bruteforce) on a classical CPU, without using the quantum annealer or any heuristic solver. The results therefore reflect our QUBO formulation alone, free of solver limitations such as noise, and test whether its optimal bipartitions yield exact species trees.

The average RF rates (over 20 replicates) of QUANTA, wQFM, wQMC and ASTRAL on varying numbers of estimated and true gene trees on the 11-taxon dataset are shown in Figure 1(a, b). Across all model conditions, these methods achieved comparable accuracy, with no statistically significant differences between any pair of methods (*p* ≥ 0.05).

On the 15-taxon dataset, we evaluated performance under varying gene tree estimation error, using 100 and 1000 bp sequence lengths, and varying numbers of genes (100 and 1000), as shown in Figure 1(c, d). QUANTA and wQFM achieved comparable accuracy, and both had lower average RF rates than ASTRAL and wQMC in all model conditions with estimated gene trees, and the improvements were statistically significant with 1000 genes and 100-bp sequences (*p* = 0.016). All methods became more accurate as the number of genes and the sequence length increased, and with true gene trees all methods recovered the model tree in every replicate.

#### 37- and 48-taxon datasets

On these dataset, partitions exceed 20 taxa, so QUANTA sends the partitions with ≥ 20 taxa to the quantum annealer. We first verified that the choice of solver does not affect accuracy: solving the same QUBOs with simulated annealing and tabu search in place of the quantum annealer yielded the same species trees on 239 of the 240 replicates across different model conditions in 37-taxon dataset, and on all 114 replicates on the 48-taxon replicates. On the single remaining replicate on 37-taxon dataset, the quantum annealer was the more accurate (details in the Supplementary Section 4). The results we discuss here are therefore for QUANTA with quantum annealing.

Figure 2 shows the results on the 37-taxon dataset as we vary the amount of ILS (0.5X being the highest), the sequence length and the number of genes. QUANTA and wQFM produced trees with comparable accuracies: the same RF distance in 230 of the 240 runs. Both QUANTA and wQFM in general produced slightly better species trees than ASTRAL and wQMC, and their improvements over ASTRAL and wQMC are statistically significant on the 250bp model condition. All methods follow expected trends: the error rates decreased as ILS decreased, the sequence length increased, or the number of genes increased.

**Figure 2:**
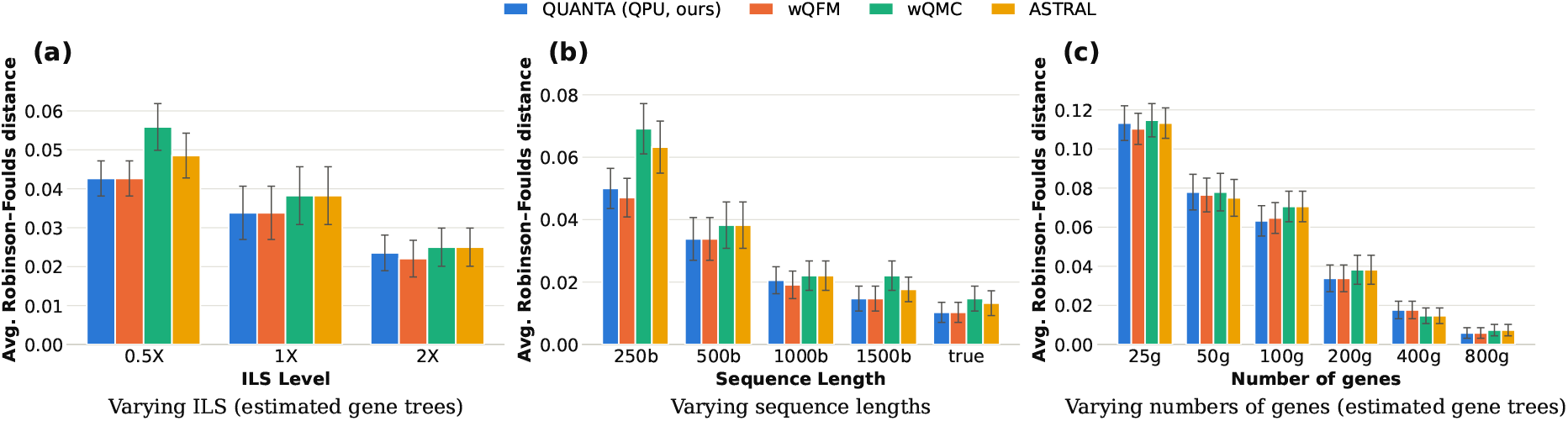
Average RF rates (20 replicates) on the 37-taxon dataset as we vary (a) the ILS level, (b) the sequence length and (c) the number of genes.

Figure 3(a, b) shows the results on the 48-taxon dataset with varying ILS levels and numbers of genes, using 10 replicates per model condition. We observed trends similar to those on the 37-taxon dataset. ASTRAL was slightly less accurate than QUANTA and wQFM, significantly so only with 200 genes (*p <* 0.05), and wQMC was the least accurate method, significantly worse than wQFM in all model conditions and than QUANTA in all but one (*p <* 0.05). All methods became more accurate as ILS decreased or the number of genes increased.

**Figure 3:**
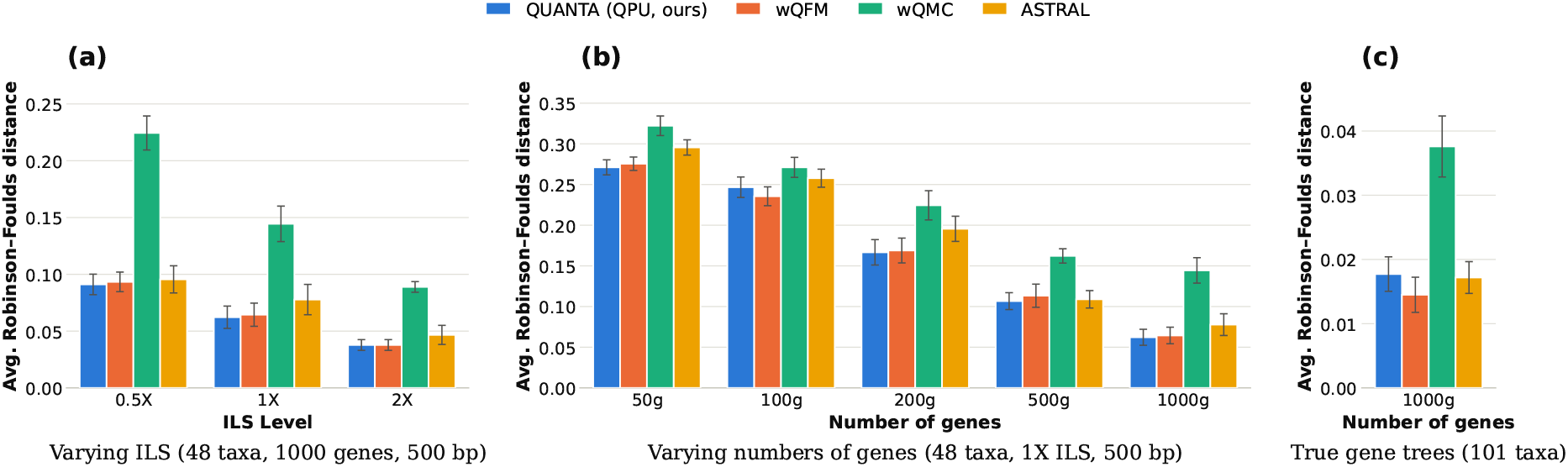
Average RF distance on the 48-taxon dataset as we vary (a) the ILS level (1000 genes, 500 bp) and (b) the number of genes (1X ILS, 500 bp), and (c) on the 101-taxon dataset (1000 true gene trees). Bars show means over 10 replicates (48 taxa) or 19 replicates (101 taxa) with standard errors.

**Figure 4:**
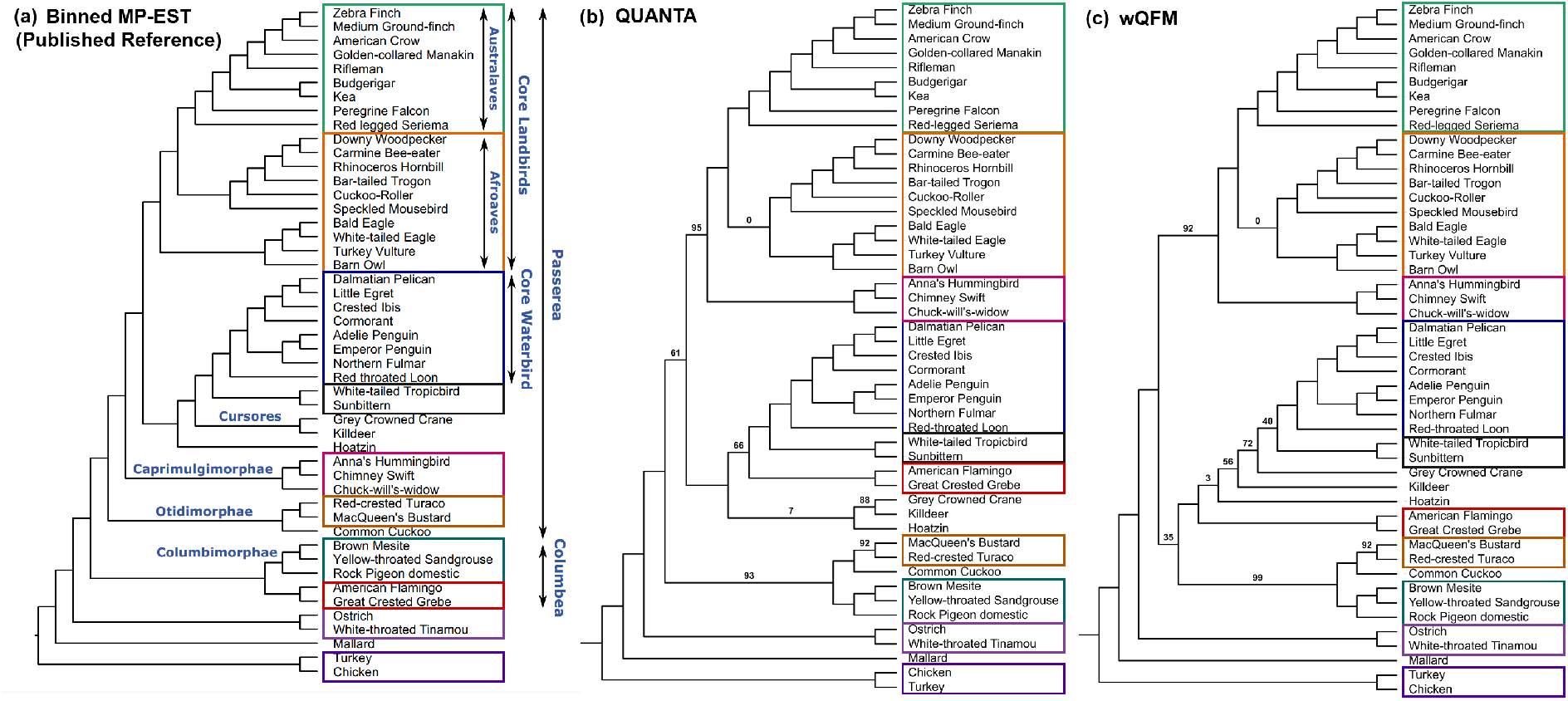
(a)-(c) Estimated trees on the avian dataset with identical clades indicated. Branch supports (BS) based on quartet-based local posterior probabilities [68] are shown on branches.

#### 101-Taxon Dataset

Results on the 101-taxon dataset with 1000 true gene trees are shown in Figure 3(c). wQFM, QUANTA, and ASTRAL produced trees with comparable accuracy with no statistically significant differences among them (*p* ≥ 0.05). Similar to other datasets, wQMC produced the least accurate trees and was significantly worse than each of the other three methods (*p <* 0.01).

#### 200-Taxon Dataset

Due to limited access to quantum hardware, we evaluated QUANTA on datasets of up to 200 taxa using real D-Wave quantum annealers and conducted additional scalability experiments on larger datasets (500 and 1,000 taxa) using simulated annealing, as described in the following section. For the 200-taxon experiment, we selected a gene tree from the ASTRAL-II simulation study [57] and extracted all of its approximately 64.7 million quartets as input. Since every quartet comes from the same tree, the correct answer is known: the tree itself, which agrees with all of them. The top-level partition of 200 taxa is too large to fit directly on the quantum processor, so it was solved with D-Wave’s hybrid solver, while the 15 smaller partitions with more than 20 taxa ran directly on the quantum annealer. QUANTA recovered the gene tree exactly (RF distance 0), satisfying all 64.7 million quartets, and so did wQFM and wQMC. Solving the same QUBOs with simulated annealing and tabu search in place of the quantum solvers also recovered the tree exactly, showing that our formulation yields the correct tree at this scale regardless of the solver.

### 3.2 Applications to scalable tree variants quartet amalgamation methods

TREE-QMC and wQFM-TREE are scalable tree variants of wQMC and wQFM that compute quartet scores directly from input gene trees, avoiding the explicit enumeration of Θ(*n*^4^) weighted quartets. Since they retain the MaxCut and FM heuristics for bipartitioning, respectively, our generalized QUBO formulation can also be applied to these methods. To demonstrate this, we developed QUANTA-TREE by replacing the FM heuristic in wQFM-TREE with our QUBO formulation and evaluated it on the 500- and 1,000-taxon simulated datasets used in the ASTRAL-II and wQFM-TREE studies [57, 66]. Due to limited access to quantum hardware, we used a classical simulated annealer to solve the QUBO problems in these experiments. The original wQFM [28] implementation could not scale to datasets of this size, while its tree variant, wQFM-TREE [29], and ASTRAL [67] required 40.9 and 25.2 minutes, respectively, for 500 taxa, and 3.2 and 2.0 hours for 1,000 taxa (results on ASTRAL and wQFM-TREE were obtained from [29]). In contrast, QUANTA-Tree completed required only 3.4 and 10.4 minutes for 500- and 1000-taxon dataset, respectively, even with classical simulated annealing. Moreover, QUANTA-TREE maintained phylogenetic accuracy comparable to wQFM-TREE and was significantly more accurate than ASTRAL (Section 4, Figure 5 in Supplementary Material).

**Figure 5:**
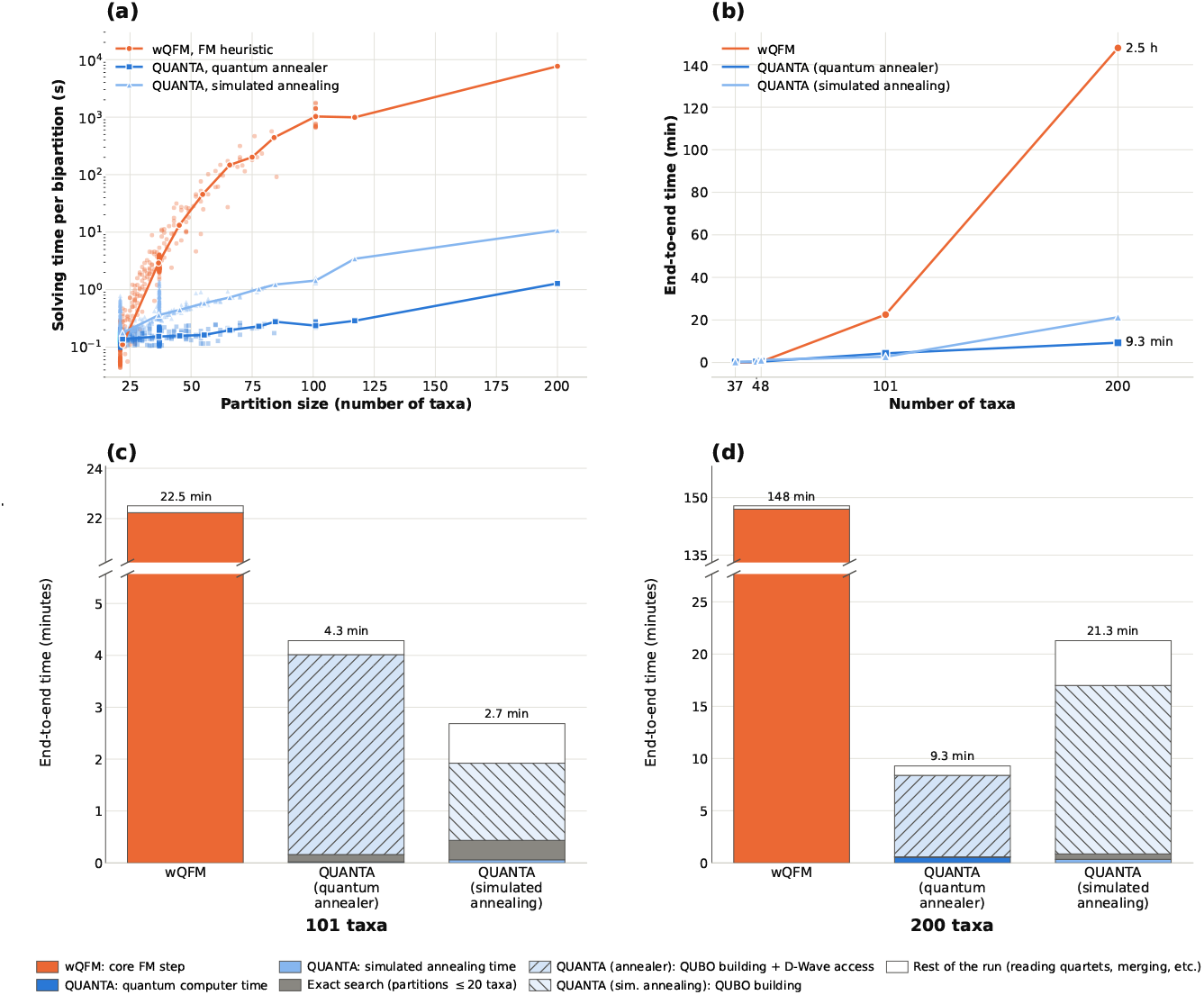
Running time of QUANTA and wQFM. (a) Solving time of each bipartition with more than 20 taxa across all datasets, for wQFM’s FM heuristic and for QUANTA with the quantum annealer (QPU access time) or simulated annealing. Points are individual bipartitions; (b) End-to-end running time by number of taxa. Each point is the mean over the replicates of one dataset. (c, d) Breakdown of the average end-to-end running time of QUANTA on 101- and 200-taxon datasets.

### 3.3 Biological Datasets

We analyzed the avian dataset comprising 48 taxa and 14,446 genes [50]. This dataset is particularly challenging due to extensive gene tree discordance, possibly resulting from rapid evolutionary radiation among ancestral lineages. Remarkably, QUANTA recovered a tree that achieved a higher quartet score (1,08,551 more agreeing quartets) than wQFM and is highly congruent with the published tree estimated by binned MP-EST analyzes. Both wQFM and QUANTA reconstructed the major clades Australaves, Afroaves, Cuprimulgimorphae, Core Waterbirds and Columbimorphae. ASTRAL, wQFM and wQMC failed to recover Cursores (crane and killdeer) [28] but QUANTA reconstructed this clade. Moreover, ASTRAL failed to recover Otidimorphae (bustard, turaco and cuckoo), whereas both wQFM and QUANTA reconstructed this clade. Both wQFM and QUANTA correctly placed seriemas as sister to the clade containing passerimorphae (passeriformes and parrot) and falcon, and thus reconstructed the well-established Australaves clade (passeriformes, parrot, falcon and seriema). However, wQMC misplaced seriemas and failed to recover Australaves [28]. All of these methods failed to recover Columbea (flamingo, grebe, pigeon, mesite and sandgrouse).

We also analyzed the mammalian and the Amniota (both DNA and AA) datasets. All methods (QUANTA, ASTRAL, wQFM, and wQMC) produced identical trees on this dataset (see Section 4, Figure 8 in Supplementary Material).

### 3.4 Running Time

We compared the running times of QUANTA and wQFM, which share the same reconstruction framework but differ in their bipartitioning strategies. QUANTA involves three main steps: (i) constructing the QUBO matrix from the input quartets, (ii) submitting the QUBO to remote quantum hardware, including mapping logical variables to physical qubits through minor embedding, and (iii) solving the QUBO on the quantum processor. To distinguish the computational advantage of quantum optimization from the overheads of accessing current quantum hardware, we compared the methods at two levels: the time spent solving bipartitions and the overall end-to-end running time. Only step (iii) replaces the search performed by FM. Step (i) is a pass over the quartets of the partition that FM also makes when scoring candidate bipartitions to find an initial bipartition, and stage (ii) is a cost of reaching remote quantum hardware rather than of the computation itself. We therefore compare running times at two levels: the time spent solving bipartitions only and the overall end-to-end wall-clock times, which include all overheads. We further examined the contribution of individual components to the total running time. To separate the contribution of the quantum hardware from that of the formulation, we also report QUANTA with simulated annealing on the same QUBOs and with the same number of samples.

#### Bipartitioning step

Figure 5(a) shows the solving time of every bipartition with more than 20 taxa across all simulated and biological runs, plotted against partition size. Lines connect the median solving times within consecutive 10-taxon bins (21–30, 31–40, etc.). The running time of FM increased sharply with partition size, from approximately 0.1 seconds for 21–25 taxa to over 1,000 seconds for around 100 taxa and more than two hours for the 200-taxon root partition. In contrast, the quantum annealer solved partitions of up to 120 taxa in approximately 0.14–0.29 seconds, with little variation across partition sizes. Consequently, the speedup over FM increased with partition size, exceeding 100-fold for partitions of 41–60 taxa and 4,100-fold for partitions larger than 90 taxa. We also evaluated classical simulated annealing on the same QUBOs. Although substantially faster than FM, simulated annealing became increasingly slower than quantum annealing as partition size increased, because the work in each of its sweeps grows with the number of interacting pairs of taxa. Quantum processor achieving up to a 12-fold speedup on partitions larger than 100 taxa. These results demonstrate the substantial computational advantage of quantum annealing for the bipartition optimization step.

#### End-to-end running time

We next compared the over-all running times, including QUBO construction, communication, embedding, and other overheads associated with accessing remote quantum hardware (Figure 5b). Despite these additional costs, QUANTA was substantially faster than wQFM on larger datasets. For 101 taxa, QUANTA completed the analyses in 4.3 minutes, compared with 22.5 minutes for wQFM, corresponding to a 5.3-fold speedup. The advantage was even more pronounced for 200 taxa, where QUANTA completed the analysis in just 9.3 minutes, compared with 2.5 hours for wQFM–a 16-fold speedup. On smaller datasets (37–48 taxa), however, the overhead of accessing remote quantum hardware out-weighed the savings in bipartition optimization, making QUANTA slightly slower than wQFM. Finally, to separate the contribution of the quantum hardware from that of the formulation, we ran QUANTA with simulated annealing, which was faster than the quantum annealer for 101 taxa (2.7 versus 4.3 minutes) but more than twice as slow for 200 taxa (21.3 versus 9.3 minutes), as its solving time grows with partition size, while still being about 7 times faster than wQFM. Overall, these results show that quantum-assisted bipartitioning can accelerate the complete species tree reconstruction pipeline, even on current noisy intermediate scale quantum hardware.

#### Running time breakdown

To understand the difference between the large speedups in bipartition solving and the more modest end-to-end improvements, we decomposed the running times into individual components (Figure 5(c, d)). For the 101- and 200-taxon datasets, quantum processing accounted for less than 1% of QUANTA’s division-step time, and exaustive search of small partitions for 0.4–3.5%, while 93–96% was spent on other costs: building the QUBOs, minor embedding and communication with the remote quantum hardware. The steps outside the division steps, mainly reading the quartet file, took 6–10% of the end-to-end time and the same absolute time as in wQFM, confirming that the two methods differ only in the division step. Thus, although quantum annealing solves bipartitions extremely quickly, the overall running time is currently dominated by classical preprocessing and hardware-access overheads. Importantly, these overheads are not intrinsic to quantum optimization and offer opportunities for further improvements, particularly through more efficient QUBO construction (for example by computing the QUBO directly from gene trees instead of enumerating quartets), reuse of embeddings by computing a single clique embedding once and reusing it for every partition, and reduced communication costs. The results therefore highlight both the substantial acceleration already achieved by QUANTA and the potential for further end-to-end improvements as quantum hardware and its supporting infrastructure advance.

## 4 Conclusions

We introduced QUANTA, the first quantum formulation of weighted quartet amalgamation and, to our knowledge, the first species tree estimation method for multi-locus data to run on real quantum hardware. On D-Wave quantum annealers, QUANTA matched the accuracy of wQFM, and was more accurate than ASTRAL and wQMC on most datasets. The quantum annealer solved bipartitions over 4100× faster than FM on partitions of more than 90 taxa. Because quantum processor (QPU) access is costly and time-limited, we evaluated QUANTA on current quantum hardware with datasets of up to 200 taxa. Although our experiments were limited to 200 taxa, we estimate from the problem-size limits in D-Wave’s solver documentation [38] that D-Wave’s hybrid quantum–classical solver can handle bipartitions of up to about 20,000 taxa with the same QUBO formulation.

Notably, performance gain QUANTA achieved over classical FM is using the current NISQ hardware. Since our formulation is proven correct independently of the hardware, it will run unchanged on future machines, such as IBM’s planned fault-tolerant computer [69] and Google’s million-qubit roadmap [70]. Beyond quartet amalgamation, our formulation offers a promising approach for extending quantum optimization to more general quartet-based species tree inference problems, such as the maximum quartet support species tree problem addressed by ASTRAL. QUANTA thus lays a foundation for quantum species tree estimation that will grow as quantum hardware matures.

## Supporting information

Supplementary material for QUANTA:

