## Supplementary material for QUANTA: for "QUANTA: A quantum annealing framework for species tree estimation using weighted quartet amalgamation"

#### 1 Satisfied, violated and deferred quartets

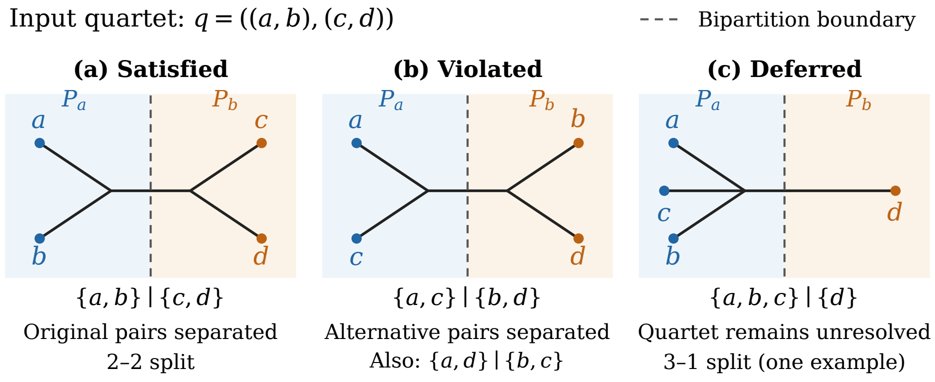


**Figure 1:** Quartet status under a bipartition: (a) satisfied, (b) violated, and (c) deferred. The dashed line marks the bipartition boundary; only the four quartet taxa are shown.

#### 2 Optional Partition-Size Penalty

The quartet objective $\mathbf{x}^\top Q\,\mathbf{x}$ does not control the sizes of the two parts. Recall that a quartet $q$ with weight $w_q$ has energy $E_q = w_q\big(A\,M_{\text{pair}} + B\,M_{\text{cross}}\big) - 4B\,w_q$, where $M_{\text{pair}}$ counts its two sister pairs that the bipartition separates and $M_{\text{cross}}$ counts its four cross pairs that it places on the same side. A quartet whose four taxa all lie on the same side has $M_{\text{pair}} = 0$ and $M_{\text{cross}} = 4$, hence energy 0, so the trivial bipartition with all taxa on one side has energy 0. In contrast, a bipartition that isolates a single taxon leaves the other three taxa of every quartet containing it on one side, each such quartet has energy $(A - 2B)\,w_q$, and the bipartition therefore has positive energy whenever $A > 2B$, as in our setting ($A = 1$, $B = 0.4$). When the quartets support no split strongly, for example under high gene-tree discordance, every non-trivial bipartition can have positive energy, and the trivial bipartition becomes the global minimum. A standard way to control part sizes in a QUBO is a quadratic penalty [1,2], which plays the role of a Lagrange multiplier [3] with a fixed coefficient:

$$\min_{\mathbf{x}\in\{0,1\}^n} \left[\mathbf{x}^\top Q\,\mathbf{x} + \gamma\Big(\sum_{i=1}^{n} x_i - S_0\Big)^2\right], \tag{1}$$

where $n = |P|$, $S_0 \in [0, n]$ is the target size of $P_b$ (and $n - S_0$ that of $P_a$), and $\gamma \geq 0$ sets the strength of the penalty.

**The penalized problem is still a QUBO.** Since $x_i^2 = x_i$,

$$\Big(\sum_{i=1}^{n} x_i - S_0\Big)^2 = \sum_{i=1}^{n}(1-2S_0)\,x_i \;+\; 2\sum_{i<j} x_i x_j \;+\; S_0^2,$$

so Equation (1) is a QUBO over the same $n$ variables with matrix $Q'_{ii} = Q_{ii}+\gamma(1-2S_0)$ and $Q'_{ij} = Q_{ij}+2\gamma$ for $i < j$; the constant $\gamma S_0^2$ does not change the minimizer. The penalty therefore adds no variables and requires no change to the solver.

**Effect on the partition sizes.** Since $\sum_i x_i = |P_b|$, the penalty depends only on the size of $P_b$: if $|P_b| = S_0 + k$, it equals $\gamma k^2$. It vanishes at the target size (for non-integer $S_0$, it is smallest at the nearest integer sizes) and grows quadratically with the deviation. Choosing $S_0 = n/2$ favours equal parts, other values of $S_0$ express a preference for unequal parts, and $\gamma = 0$ recovers the unpenalized objective. Table 1 lists the penalty for the cases of interest.

**Table 1:** Partition-size penalty $\gamma\big(|P_b| - S_0\big)^2$ for representative partitions.

| Partition | Penalty | Penalty with $S_0 = n/2$ |
|---|---|---|
| $\lvert P_b\rvert = S_0$ (target size) | 0 | 0 ($n$ even), $\gamma/4$ ($n$ odd) |
| $\lvert P_b\rvert = S_0 \pm k$ | $\gamma k^2$ | $\gamma k^2$ |
| $\lvert P_b\rvert = 1$ (singleton $P_b$) | $\gamma(1-S_0)^2$ | $\gamma(n/2-1)^2$ |
| $\lvert P_a\rvert = 1$ (singleton $P_a$) | $\gamma(n-1-S_0)^2$ | $\gamma(n/2-1)^2$ |
| $P_b = \emptyset$ | $\gamma S_0^2$ | $\gamma n^2/4$ |
| $P_a = \emptyset$ | $\gamma(n-S_0)^2$ | $\gamma n^2/4$ |

**Choosing $\gamma$.** The penalty trades off against the quartet objective. If $\gamma$ is too large, it forces near-balanced splits even when the best-supported bipartition is unbalanced, for example when an outgroup separates from the remaining taxa; if it is too small, it does not remove degenerate minima. $\gamma$ should therefore be set relative to the scale of the quartet weights, and we leave its tuning to future work.

**Use in our experiments.** We set $\gamma = 0$ throughout and handle degenerate solutions when reading the samples instead. For each partition with more than 20 taxa, the solver returns 1000 samples, and QUANTA takes the lowest-energy sample with at least two taxa on each side, skipping any lower-energy sample that does not meet this condition; exhaustive search for smaller partitions applies the same condition. If no sample qualifies, QUANTA falls back to a fixed balanced split and take the next lowest-energy sample..

### 3 How Quantum Annealing Solves the QUBO

We solve the QUBO by quantum annealing, an optimization approach based on the principles of adiabatic quantum computing [4, 5]. The Fiduccia–Mattheyses heuristic used in QFM [6] and wQFM [7] starts from an initial bipartition and improves it by moving taxa between the two parts one at a time; under our binary encoding, each such move flips one bit of the current solution. Quantum annealing works differently: through superposition, a fundamental principle of quantum mechanics, it acts on all candidate bipartitions at once rather than following a single path of moves [8].

#### 3.1 Exploring Candidate Bipartitions through Superposition

In its ideal form, quantum annealing begins in an equal superposition of all $2^n$ bitstrings: measuring this state returns each bitstring with probability $1/2^n$, i.e., a uniform distribution over all bipartitions [4]. As annealing proceeds, the QUBO objective is gradually built into the quantum state, and the amplitudes of low-cost bitstrings grow while those of high-cost bitstrings shrink (Figure 2). Measuring the final state therefore returns a low-cost bitstring with high probability.

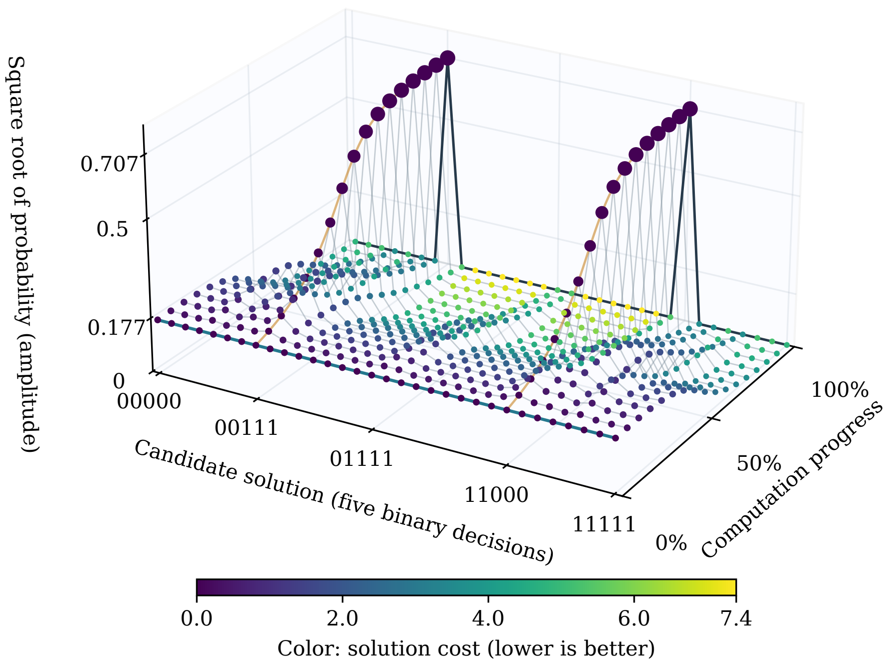


**Figure 2:** Evolution of candidate solutions during idealized five-qubit quantum annealing. Height is the amplitude (the square root of the probability of observing a bitstring) and colour is its QUBO cost. All 32 bitstrings start with amplitude $1/\sqrt{32} \approx 0.177$; at the end, the two lowest-cost bitstrings reach amplitude $1/\sqrt{2} \approx 0.707$.

#### 3.2 Time Evolution

The evolution is governed by a Hamiltonian, the operator that describes the energy of a quantum system and determines how it changes over time [4]. Quantum annealing uses the time-dependent Hamiltonian

$$H(s) = \alpha(s)\, H_{\text{driver}} + \beta(s)\, H_{\text{problem}}, \qquad s = \frac{t}{T} \in [0, 1], \tag{2}$$

where $t$ is the elapsed time, $T$ is the total annealing time, and the schedules $\alpha(s)$ and $\beta(s)$ set the weight of each term. The driver Hamiltonian has the equal superposition as its lowest-energy state and allows transitions between bitstrings [5]. The problem Hamiltonian is the QUBO objective rewritten in its equivalent Ising form, in which each binary variable becomes a two-valued spin, so that bitstrings with lower QUBO cost have lower energy [2].

During annealing, $\alpha(s)$ decreases from its maximum to zero while $\beta(s)$ increases from zero, so the energy landscape gradually changes from flat to the landscape of the QUBO (Figure 3). By the adiabatic theorem, if this change is slow enough, the system stays in its lowest-energy state, the ground state, and ends in an optimal solution of the QUBO [4, 5]. Real devices anneal for a short, finite time and are affected by noise, so the anneal is repeated many times and returns a set of low-energy samples, from which the best is taken.

#### 3.3 Escaping Local Minima through Quantum Tunnelling

A bipartition is a local minimum when every bipartition reachable by a single move has a higher cost, even though a lower-cost bipartition exists elsewhere. Reaching that better bipartition through successive moves then requires passing through higher-cost intermediate bipartitions, which form an energy barrier. A classical search can cross the barrier only by accepting these worse solutions on the way, whereas quantum tunnelling lets the system pass through the barrier without climbing over it, providing a mechanism to escape local minima [5]. Figure 4 illustrates this mechanism schematically.

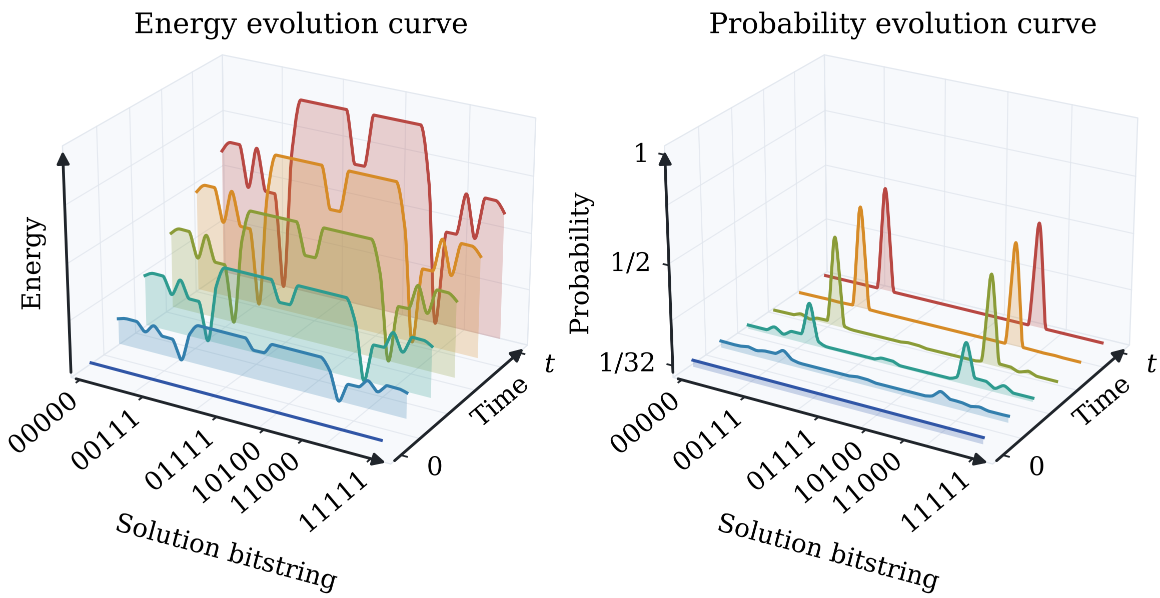


**Figure 3:** Energy and probability evolution during idealized five-qubit quantum annealing. Left: energy of each bitstring over time. Right: probability of measuring each bitstring. Initially the energy landscape is flat and all 32 bitstrings are equally likely (1/32). As annealing proceeds, 00111 and 11000 emerge as the ground states, each with final probability 1/2; they are complementary bitstrings and encode the same bipartition with its two sides swapped.

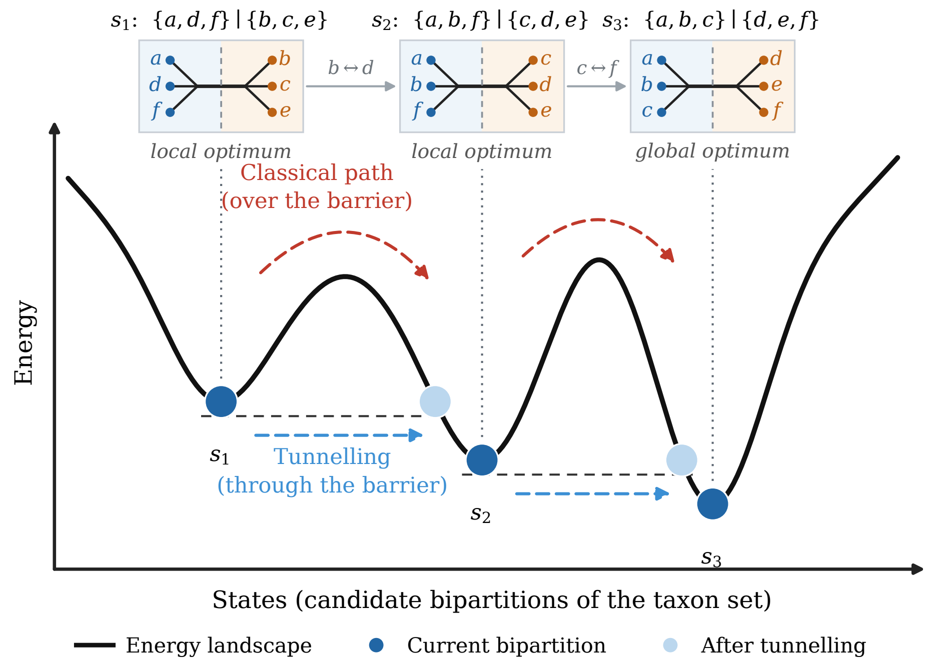


**Figure 4:** Schematic energy landscape of the bipartition search over six taxa. Lower energy indicates a better bipartition. Bipartitions $s_1$ and $s_2$ are local optima and $s_3$ is the global optimum; each step swaps two taxa between the parts ($b \leftrightarrow d$, then $c \leftrightarrow f$). A classical search must climb over each energy barrier (red, dashed), whereas quantum tunnelling passes through it (blue, dashed) toward a lower-energy bipartition.

#### 4 Additional Results

##### 4.1 Results on 500- and 1000-taxon datasets

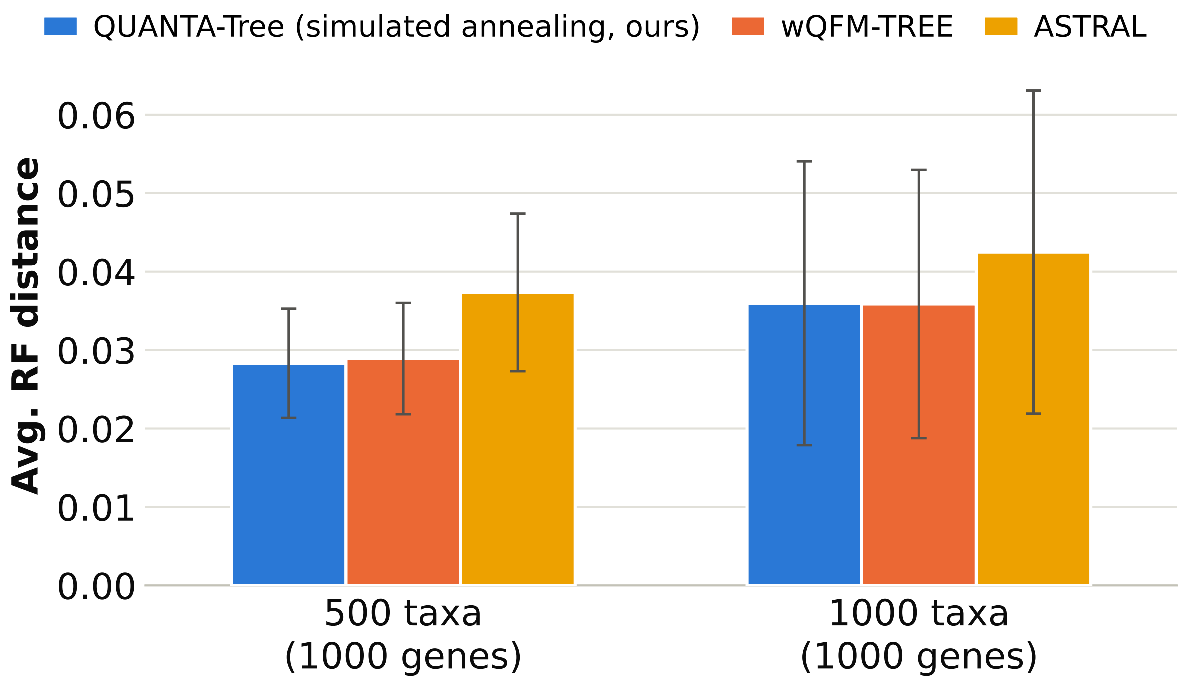


**Figure 5:** Average RF rates on the 500- and 1000-taxon datasets from ASTRAL-II study, with 1000 estimated gene trees and 10 replicates each. Error bars show the standard error. QUANTA-TREE used simulated annealing. wQFM-TREE and ASTRAL results are obtained from [9].

##### 4.2 Quantum Annealing versus Classical Solvers on the 37- and 48-Taxon Datasets

The QUBO solved at each division step depends only on the weighted quartets, not on the solver. To separate the contribution of the formulation from that of the quantum hardware, we solved the same QUBOs on the 37- and 48-taxon simulated datasets with three solvers: quantum annealing on a D-Wave quantum processor (QA), simulated annealing (SA) [10], and tabu search [11].

**Setup.** Only the solver for partitions with more than 20 taxa was changed; everything else in QUANTA was identical across the three runs, including the QUBO ($A = 1$, $B = 0.4$), exhaustive search for partitions with at most 20 taxa, and the rule for selecting a solution (the lowest-energy sample with at least two taxa on each side). QA used D-Wave Advantage processors (`Advantage2_system1` for the 48-taxon dataset) with automatic minor embedding, an annealing time of 20 $\mu$s and 1000 samples per partition. SA and tabu search used the open-source implementations in D-Wave's `dwave-samplers` package (v1.8.0) on a laptop CPU (Intel Core i5-1335U), also with 1000 samples per partition: SA performed its default 1000 sweeps per sample, and tabu search ran with its default time limit of 20 ms per sample. The 37-taxon mammalian dataset comprises 240 inputs (12 model conditions with 20 replicates each) and the 48-taxon avian dataset 114 inputs (7 model conditions). Accuracy is the normalized Robinson–Foulds (RF) distance to the true species tree.

**Accuracy.** The three solvers produced the same species trees (Figures 6 and 7). On the 48-taxon dataset, QA, SA and tabu search returned identical tree topologies on all 114 inputs, with a mean RF distance of 0.1388. On the 37-taxon dataset, they returned identical topologies on 239 of the 240 inputs (mean RF 0.0395 for QA and 0.0396 for SA and tabu search). On the remaining input (200 genes, 1000 bp, replicate 6), QA recovered the true species tree exactly, whereas SA and tabu search, which agreed with each other, placed one of the 34 internal branches incorrectly (RF $1/34 \approx 0.029$). SA and tabu search agreed with each other on all 354 inputs, and neither ever needed the fallback split. The accuracy of QUANTA is therefore determined by the QUBO formulation: whichever solver minimizes it, the same bipartitions, and hence the same trees, are found.

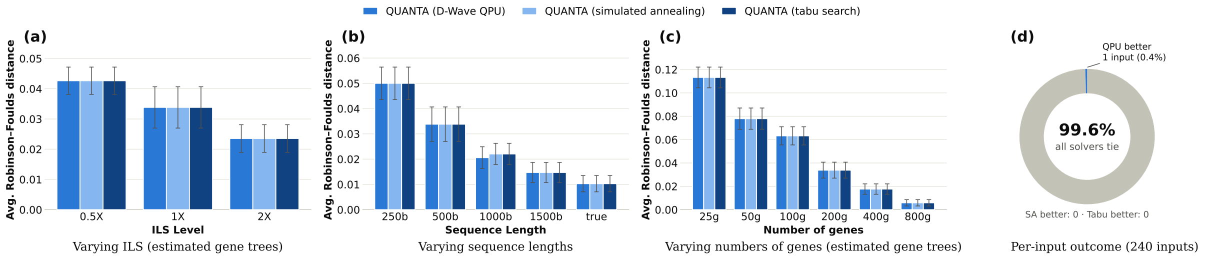


**Figure 6:** QUANTA on the 37-taxon dataset with three QUBO solvers for partitions with more than 20 taxa: D-Wave quantum annealing (QPU), simulated annealing and tabu search. (a)–(c) Average RF distance to the true species tree, with standard-error bars over 20 replicates, for varying ILS levels, sequence lengths and numbers of genes. (d) Per-input outcome: all three solvers give the same RF distance on 239 of the 240 inputs (99.6%); on the remaining input, the QPU gives the lower RF distance.

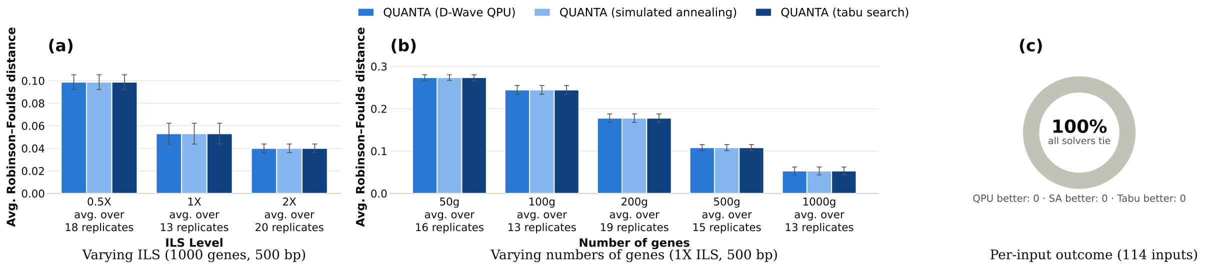


**Figure 7:** QUANTA on the 48-taxon dataset with three QUBO solvers for partitions with more than 20 taxa: D-Wave quantum annealing (QPU), simulated annealing and tabu search. (a)–(b) Average RF distance to the true species tree, with standard-error bars, for varying ILS levels and numbers of genes; the number of replicates per condition is shown under each tick. (c) Per-input outcome: all three solvers give the same RF distance on all 114 inputs.

##### 4.3 Biological datasets: Amniota and Mammalian

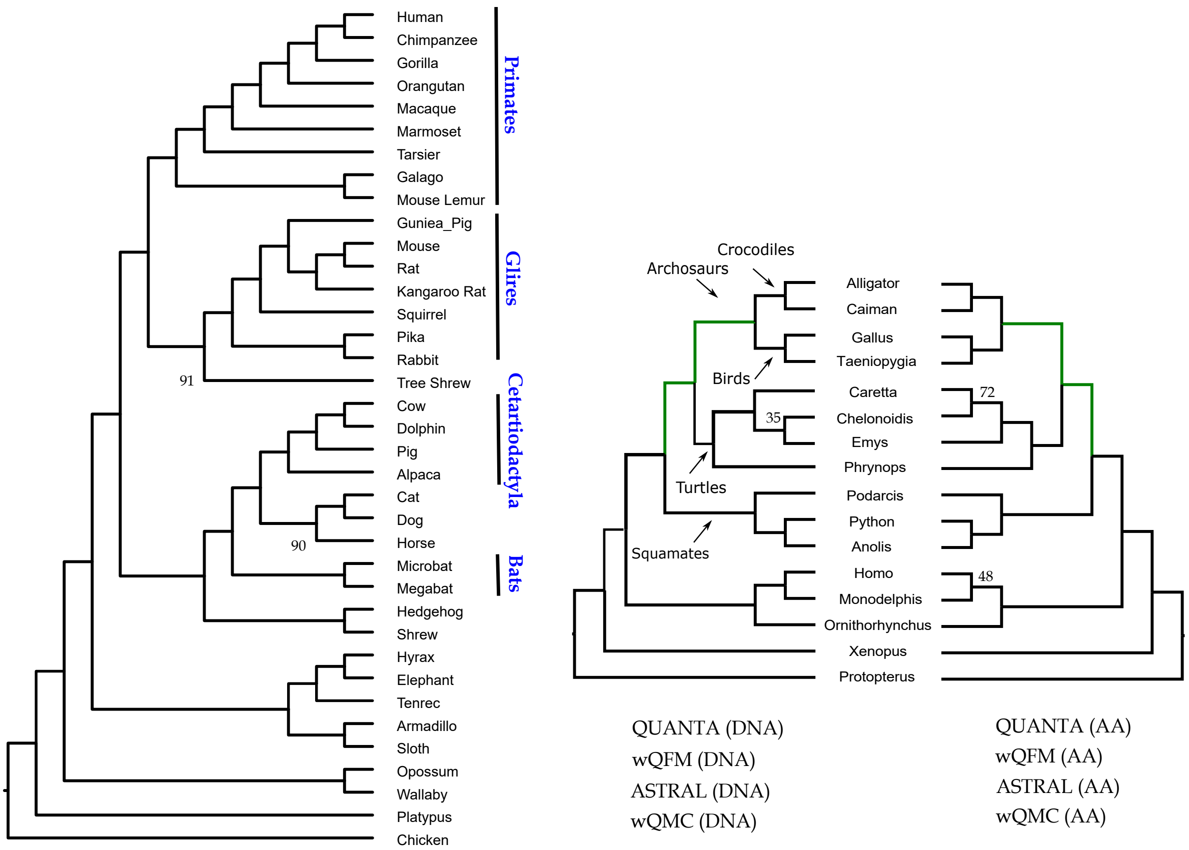


(a) Mammalian tree

(b) Amniota (DNA and AA)

**Figure 8:** Estimated trees on (a) the mammalian dataset [12], and (b) the amniota dataset [13]. Branch supports (BS) based on quartet-based local posterior probabilities [14] are shown on branches. All BS values are 100% except where noted.

#### 5 Problem-size limits of the D-Wave solvers

In QUANTA, each taxon of a partition is one QUBO variable, that is, one logical qubit. Every pair of taxa appears together in at least one quartet, so every pair of logical qubits is linked by a quadratic term, and a partition of $n$ taxa gives a dense QUBO with $n(n-1)/2$ quadratic terms. The largest such QUBO that a solver can handle depends on the solver. We assigned each partition to exhaustive search, the quantum processor (QPU) or D-Wave's hybrid quantum–classical solver according to its size (Table 2).

**Table 2:** Solvers used for the bipartition QUBO, the partition sizes assigned to each, and their estimated size limits for dense QUBOs.

| Solver | Partitions solved | Estimated limit (taxa) | Basis of the estimate |
|---|---|---|---|
| D-Wave QPU | 21–150 taxa | 175–177 (180–184 defect-free) | clique embedding |
| D-Wave hybrid solver | $>150$ taxa | about 20,000 | documented solver limits |

##### 5.1 Direct solution on the quantum processor

A D-Wave QPU consists of physical qubits connected by couplers, and each physical qubit is coupled to only 15 others on Advantage processors (Pegasus topology [15]) and 20 others on Advantage2 processors

(Zephyr topology [16]). A QUBO is solved on the QPU by minor embedding [17]: each logical qubit is represented by a chain of physical qubits that are strongly coupled so that they act as one, and each quadratic term is placed on a coupler between two chains. In a dense QUBO every chain must touch every other chain, so chains grow longer as the number of taxa increases, and beyond a certain size no embedding exists.

We estimated this size with the clique-embedding routine of D-Wave's `minorminer` package (v0.2.22) [18], which embeds fully connected QUBOs with short chains. We applied it to defect-free versions of the Advantage qubit graph (Pegasus, 5,640 physical qubits) and the Advantage2 qubit graph (Zephyr, 4,800 physical qubits), generated with `dwave-networkx` (v0.8.19), and searched by bisection for the largest number of logical qubits that could be embedded. The largest dense QUBO that fits has 180 logical qubits on Advantage and 184 on Advantage2. These embeddings use only 53% and 50% of the physical qubits, so the limit is set by the sparse connectivity of the processor rather than by its number of qubits. Real processors also have some inactive qubits and couplers, and previous studies report dense QUBOs of up to 175–177 logical qubits on Advantage processors [19, 20].

In our analyses, the QUBOs were embedded by D-Wave's automatic minor-embedding tool [21], which works for any QUBO but can produce longer chains than the dedicated clique embedding. We therefore sent to the QPU only partitions of up to 150 taxa, a conservative threshold below the limits above. The largest partition solved on the QPU had 117 taxa and came from the 200-taxon dataset.

##### 5.2 Hybrid quantum–classical solver

D-Wave's hybrid solver for binary quadratic models combines classical heuristics with calls to the QPU, so its problem size is not limited by embedding. It accepts at most $10^6$ variables and $2 \times 10^8$ linear and quadratic terms [22]. A dense QUBO of $n$ taxa has $n$ linear and $n(n-1)/2$ quadratic terms, $n(n+1)/2$ in total, so the term limit is reached first, at $n \approx 20{,}000$ taxa. The solver also requires a minimum run time that grows with the number of variables: interpolated linearly from D-Wave's documented values, it is 3 s up to 1,024 variables, about 30 s at 8,000 and about 120 s at 20,000. These are capacity limits: the solver accepts problems of these sizes, but we did not test its solution quality beyond the 200-taxon partition solved in this study.

##### 5.3 Exhaustive search

Exhaustive search evaluates all $2^n$ assignments of a partition and therefore always returns the optimal bipartition. In our implementation, a 20-taxon partition took 6.6–7.4 s (median 6.8 s over 136 partitions), and each additional taxon doubles this time. We therefore solved partitions of at most 20 taxa exhaustively, which also saved QPU access time.

#### References


[1] Fred Glover, Gary Kochenberger, and Yu Du. A tutorial on formulating and using qubo models. arXiv preprint arXiv:1811.11538, 2018.

[2] Andrew Lucas. Ising formulations of many np problems. Frontiers in physics, 2:5, 2014.

[3] Lieven Vandenberghe and Stephen Boyd. Convex optimization, volume 1. Cambridge university press Cambridge, 2004.

[15] Kelly Boothby, Paul Bunyk, Jack Raymond, and Aidan Roy. Next-generation topology of d-wave quantum processors. arXiv preprint arXiv:2003.00133, 2020.

[16] D-Wave Quantum Inc. Topologies. D-Wave Documentation, 2026. Accessed 2 November 2026.

[17] Vicky Choi. Minor-embedding in adiabatic quantum computation: I. the parameter setting problem. Quantum Information Processing, 7(5):193–209, 2008.

[18] Tomas Boothby, Andrew D King, and Aidan Roy. Fast clique minor generation in chimera qubit connectivity graphs. Quantum Information Processing, 15(1):495–508, 2016.

[19] Elijah Pelofske, Georg Hahn, and Hristo N Djidjev. Noise dynamics of quantum annealers: estimating the effective noise using idle qubits. Quantum Science and Technology, 8(3):035005, 2023.

[20] Myeonghwan Seong and Daniel Kyungdeock Park. Hamiltonian formulations of centroid-based clustering. Frontiers in Physics, 13:1544623, 2025.

[21] Jun Cai, William G Macready, and Aidan Roy. A practical heuristic for finding graph minors. arXiv preprint arXiv:1406.2741, 2014.

[22] D-Wave Quantum Inc. D-Wave quantum. `https://www.dwavequantum.com`, 2026. Accessed: 2026-10-16.
